# Of hands, tools, and exploding dots: How different action states and effects separate visuomotor memories

**DOI:** 10.1101/548602

**Authors:** Raphael Schween, Lisa Langsdorf, Jordan A Taylor, Mathias Hegele

**Author notes:** Raphael Schween, Justus-Liebig-University, Giessen, Germany, Department of Sport Science, Kugelberg 62, D-35395 Giessen, Germany.

## Abstract

Humans can operate a variety of modern tools, which are often associated with different visuomotor transformations. Studies investigating this ability have repeatedly found that the simultaneous acquisition of different transformations appears inextricably tied to distinct states associated with movement, such as different postures or action plans, whereas abstract contextual associations can be leveraged by explicit aiming strategies. It still remains unclear how different transformations are remembered implicitly when target postures are similar. We investigated if features of planning to manipulate a visual tool, such as its visual identity or the intended effect enable implicit learning of opposing visuomotor rotations. Both cues only affected implicit aftereffects indirectly through generalization around explicit strategies. In contrast, practicing transformations with different hands resulted in separate aftereffects. It appears that different (intended) body states are necessary to separate aftereffects, supporting the idea that underlying implicit adaptation is limited to the recalibration of a body model.

## Introduction

A hallmark of human motor skill is that we can manipulate a variety of different objects and tools. The apparent ease with which we switch between skilled manipulation of different tools requires that our motor system maintains representations of different sensorimotor transformations associated with them and retrieves these based on context (Wolpert and Kawato 1998; Higuchi et al. 2007). For this to work, the brain is assumed to rely on contextual cues, i.e. sensations that allow the identification of the current context in a predictive manner (though see Lonini and colleagues (Lonini et al. 2009)). This capacity has been investigated in dual adaptation experiments, where different cues are linked with different – often conflicting – sensorimotor transformations to determine the extent with which the cues enable the formation of separate visuomotor memories (Ghahramani and Wolpert 1997; Seidler et al. 2001; Imamizu et al. 2003; Osu et al. 2004; Bock et al. 2005; Woolley et al. 2007, 2011; Hinder et al. 2008; Hegele and Heuer 2010; Howard et al. 2012, 2013, 2015; Ayala et al. 2015; van Dam and Ernst 2015; Sheahan et al. 2016, 2018; Heald et al. 2018). Whereas earlier studies have predominantly found that the posture or initial state of the body act as sufficient cues (Gandolfo et al. 1996; Ghahramani and Wolpert 1997; Seidler et al. 2001; Howard et al. 2013), a number of more recent studies observed that distinct movement plans effectively separate memories for sensorimotor transformations (Hirashima and Nozaki 2012; Howard et al. 2015; Day et al. 2016; Sheahan et al. 2016; McDougle et al. 2017; Schween et al. 2018), even when these plans are not ultimately executed (Sheahan et al. 2016).

As suggested by Sheahan and colleagues (Sheahan et al. 2016), these findings can be unified under a dynamical systems perspective of neural activity (Churchland et al. 2012). Under this framework, neural states during movement execution are largely determined by a preparatory state prior to movement onset. Assuming that distinct states of the body as well as intended movements set distinct preparatory states, errors experienced during execution could therefore be associated with distinct neural states, thus establishing separate memories for novel transformations. While this idea is supported by recent findings showing that future motor plans can serve as effective cues for the separation of newly formed sensorimotor memories, these cues still pertain to intended states of the body such as visually observable movement outcomes or final postures. In contrast, context cues that are not directly related to the state of the body appear to either not allow for the development of separate motor memories (Gandolfo et al. 1996; Howard et al. 2013) or do so only when participants become aware of their predictive power and develop distinct explicit movement plans in response to them (Hegele and Heuer 2010; van Dam and Ernst 2015; Schween et al. 2018).

In the use of modern electronic tools such as video game controllers or remotely controlled vehicles, however, there are many instances where similar postures or bodily states in general are associated with different sensorimotor transformations. These differ, however, with respect to the visual representation of tools that are controlled via bodily movements (as in the case of operating a drone or steering a virtual car by the same remote control) and with respect to the action effects that one strives to achieve. Thus, distinct preparatory states most likely incorporate features beyond bodily states such as the identity of the tool which is operated and/or the nature of the intended action effect.

Here, we considered the identity of the tool being controlled and the intended action effect as parts of a movement’s plan (i.e. its preparatory activity) and tested whether these cues would allow for the development of separate motor memories. Based on a previous study by Howard and colleagues (Howard et al. 2013), which showed that the visual orientation of a controlled object was modestly successful in separating motor memories, we expected the visually perceived identity of a tool to constitute a relevant contextual cue in establishing separate motor memories. To test this, participants in our first experiment practiced two opposing cursor rotations associated with different cursor icons or “tools”.

Our second experiment was inspired by ideomotor theory, according to which actions are represented by their perceivable effects (see Stock and Stock (Stock and Stock 2004) for a review of its history). More specifically, our approach is based on a strong version of ideomotor theory claiming that effect anticipations directly trigger actions (Shin et al. 2010). According to the theory of event coding (Hommel et al. 2001), effect anticipations are not limited to spatial properties, but can refer to any remote or distal sensory consequences anticipated in response to an action. The direct-activation hypothesis has received empirical support from neurophysiological studies showing that the mere perception of stimuli that had been established as action effects during a preceding practice phase were able to elicit neural activity in motor areas (Elsner et al. 2002; Melcher et al. 2008; Paulus et al. 2012). If we allow effect anticipations to be part of a neural preparatory state under the above view, this suggests that distinct action effects that a learner intends to achieve should allow distinct sensorimotor transformations to be associated with them. If confirmed, this would extend the state space relevant for the separation of motor memory from physical to psychological dimensions (Shepard 1987; Tenenbaum and Griffiths 2001) and thereby potentially explain separation of memory for movements with similar body states. To test this, we investigated how participants adapted to two opposing visuomotor cursor rotations when these were associated with different action effects.

Finally, we conducted a control experiment where we tested if the use of separate hands and thus clearly distinguishable bodily states would cue distinct motor memories of the opposing visuomotor transformations. Given that different effectors can be considered different states of the body and that intermanual transfer of adaptation is limited (Malfait and Ostry 2004; Sarwary et al. 2015; Poh et al. 2016), we hypothesized that this would lead to clearly separate memories for the two transformations.

## Results

### Experiment 1: Visual tools

The goal of experiment 1 was to determine if different visual tools in separate workspaces could afford dual adaptation to conflicting sensorimotor transformations. In alternating blocks, participants practiced overcoming two opposing 45° visuomotor rotations by controlling two different visual tools, either a cartoon of a hand or an arrow cursor, in separate regions of the visual workspace (figure 1A, figure 2A). The clockwise and counterclockwise perturbations were uniquely associated with either the hand or arrow cursor (and workspaces), counterbalanced across participants. To distinguish whether separate memories were formed and retrieved based on the context cue or separate explicit motor plans (Schween et al. 2018), we tested spatial generalization of learning under each cue differentially and dissociated total learning into explicit plans and implicit adaptation by a series of posttests (Heuer and Hegele 2008).

**Figure 1:**
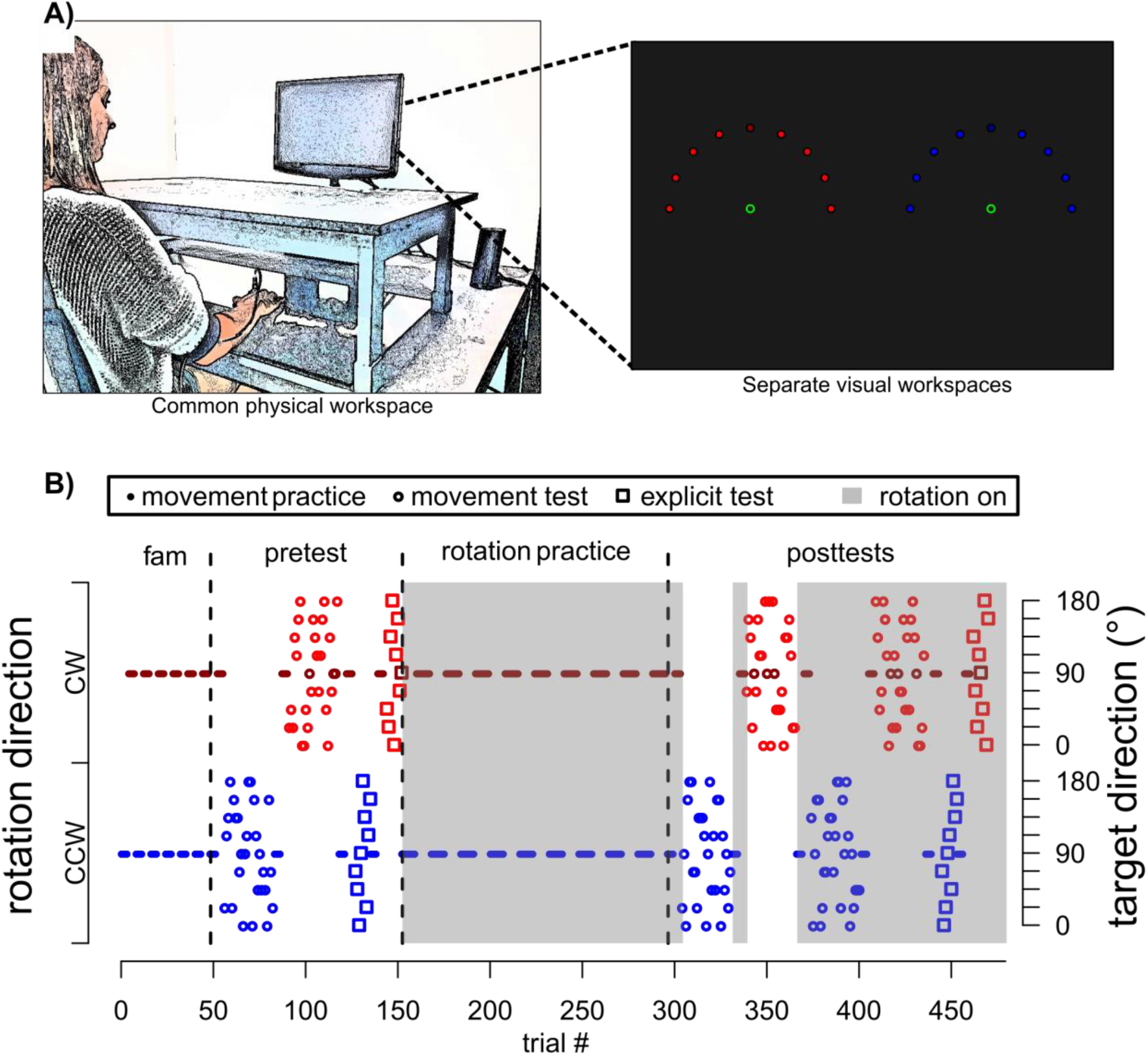
Experimental setup (A) and an exemplary task protocol (B). Adapted from Schween et al. (Schween et al. 2018) under CC BY-4.0 license. Participants made rapid shooting movements, dragging a motion-captured sled attached to their index finger across a glass surface, to bring the different tools to a single target (indicated by darker red/blue color), which was oriented at 90° and presented on a vertically-mounted computer screen. Continuous, online feedback was provided during reach practice, where cursor movement was veridical to hand movement in familiarization, but rotated 45° during rotation practice, with rotation sign and contextual cue level alternating jointly in blocks of 8 trials. Pre and posttests without visual feedback tested generalization to different targets (practice target + targets indicated in lighter red/blue) with participants either instructed that the rotation was present (total learning) or absent (aftereffects) or judging the required movement direction using a visual support (explicit judgment), from which we inferred total, implicit and explicit learning, respectively.

**Figure 2:**
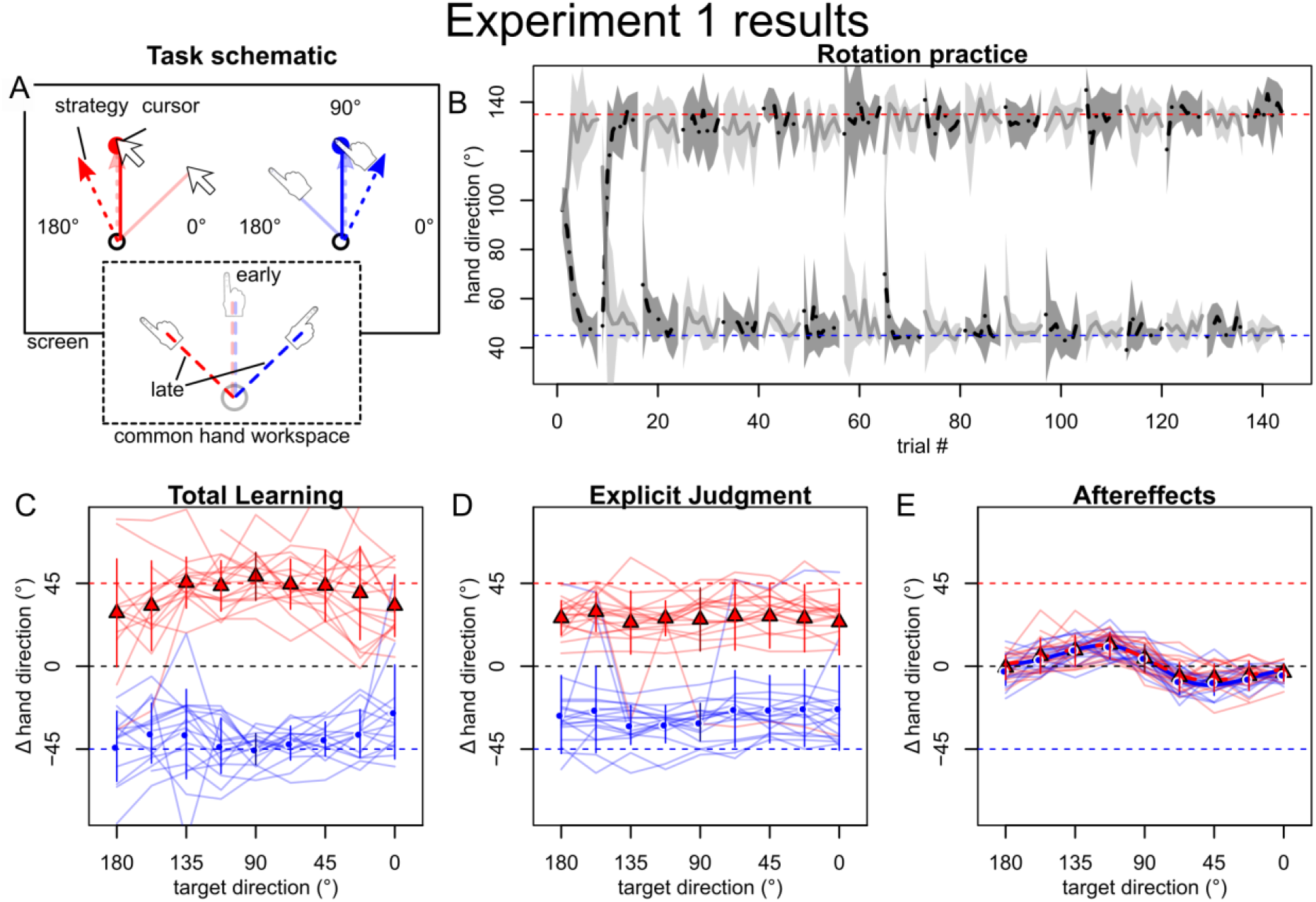
Results of experiment 1. A: Opposing rotations were cued by different screen cursors and display in different parts of the visual workspace (VWS). B: Light and dark grey lines and shades represent means and standard deviations of hand direction averaged over participants who began practice with CW rotation/left VWS or CCW rotation/right VWS, respectively (as this was counterbalanced). Participants quickly learned to compensate for both rotations, as indicated by grey lines approaching red and blue dashed horizontal lines marking perfect compensation. C-E: Symbols with error bars represent across participant means and standard deviations of baseline-corrected posttest directions by generalization target (x-axis) and rotation sign/VWS (red vs. blue). Note that the pairing of cue level (i.e. cursor type) to rotation sign/VWS was counterbalanced across participants and the rotation-specific red and blue curves therefore contain both cue levels, but are distinct in VWS and rotation sign experienced. Thin, colored lines are individual participant values. Total learning (C) and explicit judgments (D) appeared to depend on the context cue, but implicit learning (E) only varied with test direction, and was fit better by a bimodal than a unimodal Gaussian (thick red and green lines), in line with local generalization around separate explicit movement plans.

Within a few blocks of practice, all participants were able to compensate for the opposing rotations (figure 2B). Total learning, which is believed to comprise both explicit and implicit contributions, generalized broadly across the workspace in a pattern that is suggestive of dual adaptation (figure 2C). Much of the dual adaptation could be attributed to different explicit judgments (figure 2D) for each tool, while implicit aftereffects appeared to contribute very little to the total learning (figure 2E). What’s more, the pattern of generalization of the aftereffects appeared to be similar for each tool and exhibited a bimodal shape, indicative of plan-based generalization (figure 2E). These findings are remarkably similar to our previous findings, which had just visual workspace separation as contextual cue (Schween et al. 2018), suggesting that distinct visual tools did not afford greater dual adaptation.

As our primary goal was to determine if distinct tools could serve as cues to separate implicit visuomotor memories, we sought to further characterize the pattern of interference (or generalization) of the aftereffects. Here, we found that the mean aftereffects across generalization targets were fit better by the sum of two Gaussians (“bimodal” Gaussian) than a single Gaussian (ΔBIC: 16.1 for cued CW rotation, 17.0 for cued CCW rotation), which is consistent with our previous findings (Schween et al. 2018). The gain parameters had opposite signs and their respective bootstrapped confidence intervals did not include zero (table 1), suggesting that adaptive changes in response to both rotations were represented in each generalization function. The locations and signs of the peaks comply with what we previously explained by implicit adaptation for each of the two practiced rotations generalizing narrowly around the cue-dependent explicit movement plans (Schween et al. 2018). Here, the bimodal curve can be thought of as the sum of these independent generalization functions, where the two modes reflect the two opposite peaks of the individual functions and interference is maximal at the practiced target. Importantly, confidence intervals of differences between bootstrapped parameters for the two curves included zero (table 1), indicating that adaptation retrieved under the two context cue levels did not differ significantly. In summary, we take these results to show that visual tools did not cue separate implicit visuomotor memories, except indirectly, mediated by plan-based generalization around separate explicit movement plans.

**Table 1:** Parameters of generalization functions fit to across-participant mean aftereffects. Only parameters for the better-fitting model (unimodal or bimodal Gaussian) are shown. Square brackets contain 95% confidence intervals obtained by bootstrap resampling of participants and fitting to mean data. Differences between CW and CCW were calculated within each bootstrap sample to test for differences between aftereffects obtained under the cue levels. Abbreviations: Std. dev.: standard deviation. VWS: visual workspace.

| Experiment | Cued rotation | Parameters |  |  |  |  |
| --- | --- | --- | --- | --- | --- | --- |
|  |  | „gain“ a1 | „center“ b1 | „gain“ a2 | „center“ b2 | „std.dev.“ c |
| 1: tool + VWS | CW | 16.3° [10.5°; 85.1°] | 105.5° [86.7°; 124.5°] | -13.2° [-83.3°; -5.6°] | 66.0° [38.8°; 86.9°] | 47.2° [31.7°; 56.0°] |
|  | CCW | 16.3° [10.4°; 98.1°] | 103.5° [82.7°; 120.5°] | -15.4° [-97.4°; -8.2°] | 62.5° [39.3°; 84.0°] | 45.3° [33.7°; 52.4°] |
|  | CW - CCW | [-79.2°; 66.7°] | [-28.4°; 35.4°] | [-65.8°; 81.9°] | [-37.5°; 38.2°] | [-13.6°; 13.7°] |
| 2: effect + VWS | CW | 10.2° [7.6°; 48.7°] | 125.0° [94.9°; 133.4°] | -8.3° [-48.1°; -5.7°] | 44.2° [35.0°; 82.2°] | 40.4° [33.3°; 54.0°] |
|  | CCW | 13.4° [10.9°; 90.2°] | 119.3° [91.6°; 128.3°] | -9.3° [-87.4°; -5.7°] | 58.4° [36.5°; 91.8°] | 42.3° [32.0°; 53.8°] |
|  | CW - CCW | [-77.7°; 3.7°] | [-15.2°; 35.0°] | [-6.3°; 77.4°] | [-48.0°; 19.4°] | [-14.1°; 11.0°] |
| 3: hands + VWS | CW | 14.2° [11.4°; 17.5°] | 108.6° [97.8°; 118.1°] |  |  | 52.9° [44.2°; 61.8°] |
|  | CCW | -16.5° [-20.6°; -13.1°] | 75.2° [67.0°; 85.2°] |  |  | 68.2° [52.8°; 86.2°] |

### Experiment 2: Action effects

Motivated by ideomotor theory (Shin et al. 2010), experiment 2 tested whether opposing transformations could be learned and retrieved separately when they were associated with different intended effects of the motor action. For this purpose, participants were instructed that they should either “paint” or “explode” the target. The current effect was announced by an on-screen message at the beginning of each block and participants saw an animation of a brushstroke (paint-cue) or an explosion (explode-cue) where their cursor crossed the specified target amplitude and heard respective sounds. Again, we retained separate visual workspaces as in our previous experiments.

The results show no relevant qualitative differences compared to the combination of visual tool and workspace used in experiment 1. Participants quickly compensated for both rotations during practice (figure 3B). Total learning and explicit judgments compensated for the rotations in a cue-dependent fashion and generalized broadly (figure 3C-D). Aftereffects (figure 3E) displayed a bimodal pattern (ΔBIC CW: 19.7, CCW: 18.8) that is visually similar to that of experiment 1. The oppositely signed peaks again complied with plan-based generalization and bootstrapped parameters indicated no difference between the curves for the different cue levels (table 1). It therefore appears that separate action effects were ineffective in cuing separate memories for the opposing transformations, except as mediated by spatially separate explicit movement plans.

**Figure 3:**
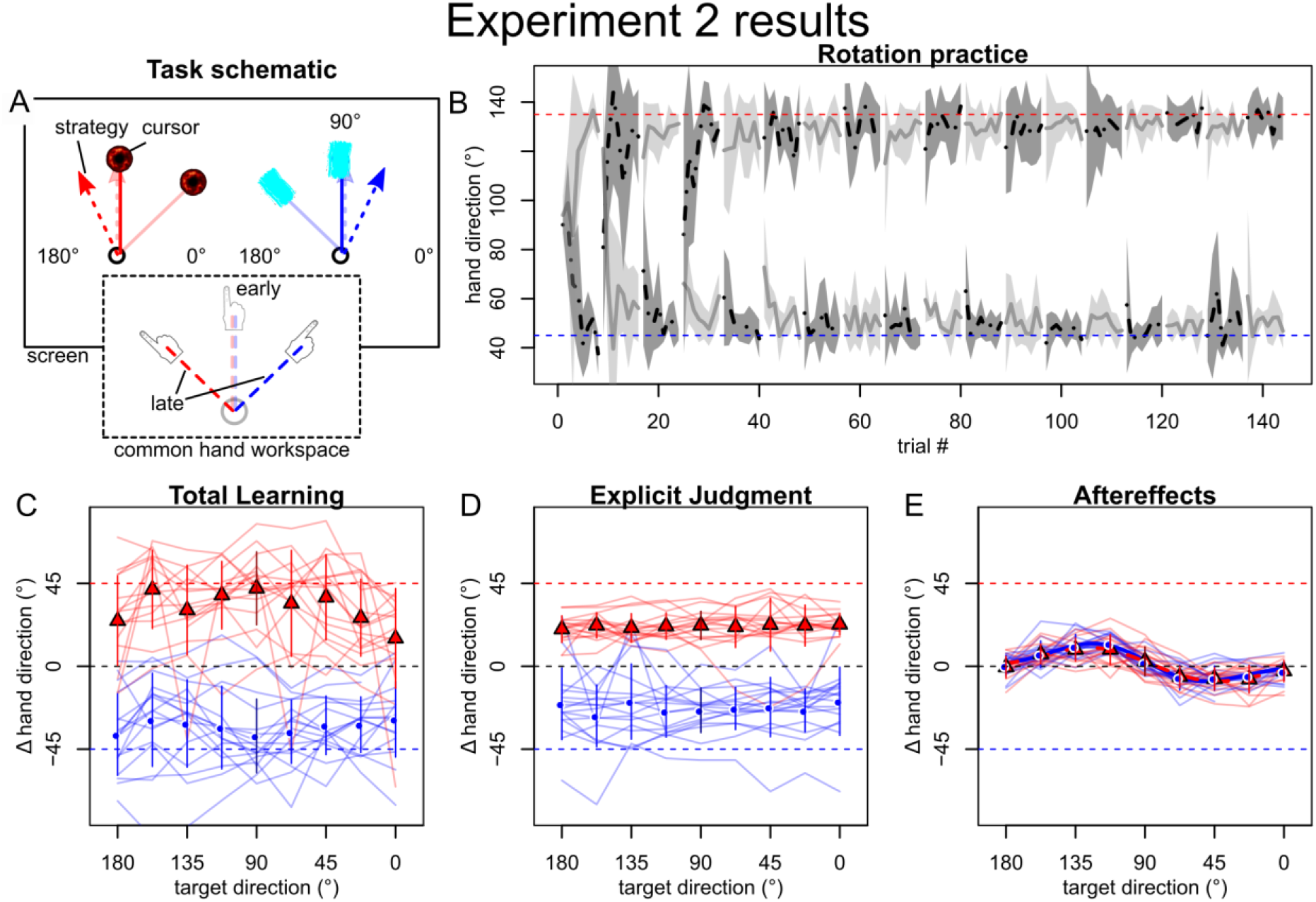
Results of experiment 2. A: Opposing rotations were now cued by participants’ intention to either “paint” or “explode” the target. Target intentions were instructed by onscreen messages at the beginning of each block and supported by animations where the cursor crossed the target amplitude, accompanied by respective sounds. B: Similar to experiment 1, mean hand directions during practice (grey lines with shades indicating SDs) indicated that participants learned to compensate both rotations, quickly. C-E: Baseline-corrected mean (±SD) and individual participant posttest directions for total learning (C) and explicit judgments (D) were specific to the contextual cue (red vs. blue) and generalized broadly across target directions (x-axis), while implicit aftereffects (E) remained cue-independent and only varied with target direction. It therefore appears that opposing transformations were learned specific to the intended effect only by explicit strategies and local, plan-based generalization around these (bimodal Gaussian fits indicated by thick red and blue lines in panel E), but no distinct implicit memories were formed depending on the intended effect.

### Experiment 3: Separate hands

As the first two context cues we tested were effective only via separate explicit strategies, we wanted to test a cue that would allow separate implicit memories to be created with relative certainty. Based on the findings that distinct body states cue separate memories and that transfer of learning between hands is incomplete, we reasoned that using different hands to practice the two rotations would be a promising approach. In experiment 3, a clockwise cursor rotation was therefore associated with left hand movements and visual display in the left half of the screen (left visual workspace), whereas a counterclockwise rotation was cued by right hand movements and right visual workspace display.

Similar to experiment 1 and 2, total learning and explicit judgments indicated that participants learned to compensate each rotation in a cue-specific fashion. Total learning at the practiced target almost completely compensated the rotation associated with each cue and relatively broad generalization to other targets occurred (figure 4C). Explicit judgments at the practiced target compensated about half of the cued rotation and also displayed a flat generalization pattern (figure 4D). Different than in the first two experiments, implicit aftereffects also showed a clear, cue dependent separation, with a single-peaked generalization pattern (figure 4E). This was supported by BIC being similar between single and bimodal Gaussian (ΔBIC CW: 1.9, CCW: 0.4, each in favor of the single Gaussian). The direction of the single Gaussians’ peaks depended on the hand cue, in line with each reflecting adaptation to the respective rotation practiced with that hand. Further, their locations were shifted off the practiced target in the direction consistent with plan-based generalization (table 1). Interestingly, generalization appeared to be considerably wider than the peaks in the first two experiments and in previous studies (McDougle et al. 2017; Poh and Taylor 2018). Furthermore, we note that, despite separate implicit memories being established, implicit learning seemed incapable of accounting for the full cursor rotation, as it was supplemented by explicit strategies, in line with recent findings that the extent of implicit adaptation is limited (Kim et al. 2018).

**Figure 4:**
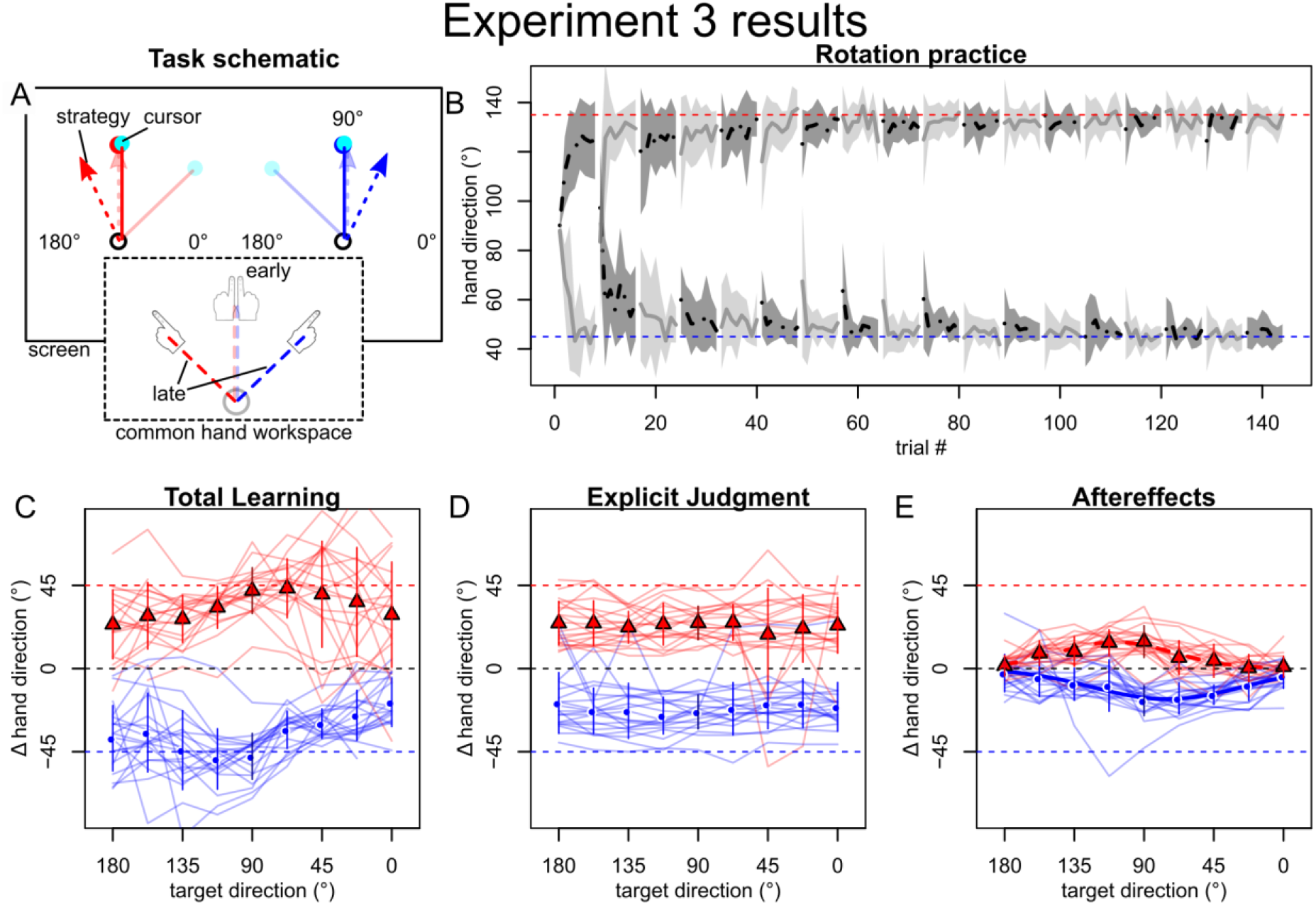
When using distinct hands to learn opposing transformations (A), participants also quickly compensated the rotation, as indicated by mean hand directions during practice (grey lines with SD shades) quickly approaching ideal compensation (red/green, dashed lines). C-E: In contrast to experiment 1 and 2, across participant mean (±SD) directions now appeared specific to the cue level (red vs. blue symbols and thin lines) for aftereffects (E) in addition to total learning (C) and explicit judgments (D) and were best fit by a unimodal Gaussian (thick red and blue lines in E).

In summary, these results indicate that the cue combination of using separate hands in addition to the separate visual workspaces successfully cued separate visuomotor memories for both implicit adaptation and explicit strategies. As contextual separation of memories can be considered the inverse of transfer between contexts, this is in line with findings suggesting that intermanual transfer of sensorimotor adaptation relies largely on strategies being flexibly applied across hand-context (Malfait and Ostry 2004; Poh et al. 2016) and that implicit learning transfers incompletely to the other hand (Sarwary et al. 2015; Poh et al. 2016) (but see Kumar and colleagues (Kumar et al. 2018)).

## Discussion

Based on the assumption that distinct preparatory states most likely incorporate features beyond bodily states, we considered the identity of a tool being controlled and the intended action effect as being part of a movement’s plan (i.e. its preparatory activity) and tested whether these cues would allow for the development of separate motor memories. We also distinguished between explicit and implicit forms of motor memories. Contrary to our expectation, neither distinct tools nor action effects appeared to produce separate implicit memories. Instead, the opposing transformations were represented in implicit memory only indirectly via local spatial generalization around explicit movement plans (Hirashima and Nozaki 2012; Day et al. 2016; Sheahan et al. 2016; McDougle et al. 2017; Schween et al. 2018). Consistent with previous findings, it appears that separate implicit memories are inextricably linked to states of the movement or the body. Indeed, in a control experiment (experiment 3), we found that distinct aftereffects, an indicator for the development of separate implicit motor memories, formed when participants practiced the opposing visuomotor rotations with separate hands, which represent a strong cue within a bodily state space. Under the dynamical systems perspective invoked in the introduction, these results would indicate that only past, current and future states of the body determine the preparatory state in areas relevant to implicit adaptation, while more abstract contextual cues that relate to the action but not to parts of the body, are processed differently.

Despite our findings, we know that people can manipulate a variety of tools with little cognitive effort, even if they share similar movement characteristics and workspaces. Thus, we would expect that humans are capable of separating and storing separate implicit memories based on cues that do not require distinct states of the body. This raises the question as to why studies have consistently failed to find contextual cues that do not depend on movement-related states (Gandolfo et al. 1996; Woolley et al. 2007; Hinder et al. 2008).

One possibility lies in the way in which context was implemented during practice and testing: it is well possible that experimental parameters like the duration of practice, the frequency and schedule of change between transformation-cue-combinations (Osu et al. 2004; Braun et al. 2009), or the way cues are presented in tests without visual feedback are responsible for the absence of implicit dual adaptation and it is a limitation of our study that we did not test different conditions. However, recent findings have shown that implicit adaptation in cursor rotation experiments approaches a fixed asymptote (Kim et al. 2018), which suggests that even longer practice would not enable this implicit adaptation process to account for large-scale change in visuomotor relations. These findings thus align with ours in suggesting that learning transformations associated with tools and contexts may rely on mechanisms distinct from this implicit adaptation.

What could be the nature of these mechanisms? Morehead and colleagues (Morehead et al. 2015) suggested that explicit movement plans are part of an action selection mechanism that can also operate implicitly, whereas implicit adaptation typically observed in cursor rotation experiments reflects a calibration mechanism for motor execution on a lower level. We speculate that the action selection level is where context-dependent learning of tool transformation occurs. Under this view, implicit, context-dependent learning could, for example, be achieved by proceduralization of strategies at the action selection level, in line with canonical views of skill learning (Fitts and Posner 1967; Willingham 1998). Recent findings have shown that that new policies can be learned by exploration and reinforcement (Shmuelof et al. 2012; Vaswani et al. 2015), and that this is closely tied to explicit strategies (Codol et al. 2018; Holland et al. 2018). A possibility is that explicit action selection tendencies may become proceduralized by associative learning. Consistent with this idea, recent findings indicate that stimulus-response associations in cursor rotation (Huberdeau et al. 2017; Leow et al. 2019; McDougle and Taylor 2019) and visuomotor association paradigms (Hardwick et al. 2018) can become “cached”, so that they are expressed by default in situations where cognitive processing resources are limited.

These roles we assign to action selection and execution are reminiscent of another canonical distinction in the motor learning literature, between the learning of body versus tool transformations (Heuer 1983; Berniker and Körding 2008, 2011; Kong et al. 2017) giving rise to modifications of an internal representation of the body (body schema) (Kluzik et al. 2008; Cardinali et al. 2009) and internal representations of tools, respectively (Massen 2013; Heuer and Hegele 2015). Here, our results would suggest that aftereffects in standard cursor rotation experiments reflect a dedicated mechanism that keeps the system calibrated to minor changes in the transformations of the body, and is therefore sensitive to context regarding the state of the body, but not other aspects of the motor environment. Notably, limiting standard implicit adaptation to a body model does not necessarily contradict the idea that internal models and the cerebellum underlie tool transformations (Imamizu et al. 2003; Higuchi et al. 2007; Imamizu and Kawato 2009). Recent neuroimaging and patient studies indicate that cerebellum-based internal models support not only the implicit (Leow et al. 2017), but also the explicit component of visuomotor adaptation (Werner et al. 2014; Butcher et al. 2017). A possibility is that internal models support the selection of suitable actions by simulating their hypothetical outcomes (Barsalou 1999).

Within the theory of event coding (TEC) (Hommel et al. 2001) mentioned in the introduction, our findings can be explained along a similar route: TEC acknowledges that neural signatures underlying perception and action need to be distinct at the far ends of this continuum and common coding and effect-based representation of actions can therefore only occur on an intermediate level (Hommel et al. 2001). As such, our results would place implicit adaptation to cursor rotations towards the action side of processing, thus explaining why separate action effects did not enable separate learning in our study, but the use of separate effectors did.

Throughout this work, we chose to call visual tools, action effects and the limb used for execution a “contextual cue”, whereas we did not consider the different targets contextual cues, that we utilized to test generalization. This reflects the arbitrary choice to define a visuomotor memory as representing the physical space of the experiment. Modelling-based accounts have assumed more clear formulations of similar ideas, e.g. in defining “contextual signals” as “all sensory information that is not an element of dynamical differential equations” (Haruno et al. 2001). However, it is unclear to which extent physical space is relevant to the neural organization of the brain and the question of how to define “contextual cue” eventually becomes the same as which contextual cues enable the formation of separate memories in which ways. In this sense, our findings mean that adaptation of the implicit body model reflected in aftereffects only responds to contextual cues that are directly related to the state of the body, whereas supposed action selection component can in principle account for any contextual cue provided that the cue is either subject to overt attention or supported by a previously reinforced association.

In conclusion, it remains a puzzle how we appear to use tools that share the same body states but require different sensorimotor transformations, most strongly exemplified in “modern” tools like video game controllers. Our work indicates that action effects and visual tools are insufficient to cue separate implicit memories for this scenario under the practice conditions studied. We speculate that, rather than implicit recalibration replacing explicit strategies with prolonged practice, these strategies may become proceduralized, with associative learning supporting the brain in acquiring the contingencies between contextual cues and appropriate action selection policies.

## Methods

Sixty-three human participants provided written, informed consent as approved by the local ethics committee of the Department of Psychology and Sport Science of Justus-Liebig-Universität Giessen and participated in the study. To be included in analysis, participants had to be between 18 and 30 years old, right handed, have normal or corrected to normal visual acuity and were not supposed to have participated in a similar experiment, before. We therefore excluded 8 participants (5 for not being clearly right-handed according to the lateral preference inventory (Büsch et al. 2009), 2 for failing to follow instructions according to post-experimental standardized questioning, one for exceeding our age limit), giving us a total of 18 analyzed participants in experiment 1, 17 in experiment 2, and 20 in experiment 3.

### Apparatus

The general task and protocol were similar to those described in our previous study (Schween et al. 2018). Participants sat at a desk, facing a screen running at 120 Hz (Samsung 2233RZ), mounted at head height, 1 m in front of them (figure 1A). They moved a plastic sled (50 × 30 mm base, 6 mm height) strapped to the index finger of their right hand (and left hand, respectively, in experiment 3), sliding over a glass surface on the table, with low friction. A second tabletop occluded vision of their hands. Sled position was tracked with a trakSTAR sensor (Model M800, Ascension technology, Burlington, VT, USA) mounted vertically above the fingertip, and visualized by a custom Matlab (2011, RRID:SCR_001622) script using the Psychophysics toolbox (RRID:SCR_002881 (Brainard 1997)), so that participants controlled a cursor (cyan filled circle, 5.6 mm diameter or specific cursors in *experiment* 1).

### Trial types

Trials began with arrows on the outline of the screen guiding participants to a starting position (red/green circle, 8 mm diameter) centrally on their midline, about 40cm in front of their chest. Here, the cursor was only visible when participants were within 3mm of the start location. After participants held the cursor in this location for 500ms, a visual target (white, filled circle, 4.8mm diameter) appeared at 80mm distance and participants had to “shoot” the cursor through the target, without making deliberate online corrections. If movement time from start to target exceeded 300ms, the trial was aborted with an error message.

Participants experienced 3 types of trials. On *movement practice* trials, they saw the cursor moving concurrently with their hand. Here, cursor feedback froze for 500ms, as soon as target amplitude was reached. On *movement test* trials, we tested behavior without visual feedback meaning that the cursor disappeared on leaving the start circle. On *explicit judgment* trials (Heuer and Hegele 2008), we asked participants to judge the direction of hand movement required for the cursor to hit the target, without performing a movement. For this purpose, participants verbally instructed the experimenter to rotate a ray that originated in the start location to point in the direction of their judgment. During judgments, they were asked to keep their hand supinated on their thigh in order to discourage them from motor imagery. Accordingly, moving towards the start position was not required.

### General task protocol

The experiment consisted of four phases: familiarization, pretests, rotation practice and posttests (figure 1B). *Familiarization* consisted of 48 movement practice trials to a target at 90° with veridical cursor feedback, thus requiring a movement “straight ahead”. The cue condition (see *Specific experiments)* alternated every 4 trials, with condition order counterbalanced across participants. *Pretests* contained movement practice tests and explicit judgment tests to establish a baseline for subsequent analysis. We tested generalization to 9 target directions from 0° to 180° at the amplitude of the practice target. We obtained one set per cue level, which in turn consisted of 3 blocks of randomly permuted trials to each of the 9 target directions for movement tests and one such block for explicit judgment tests. The sets were interspersed by blocks of 8 practice movements (4 per cue level) to the 90° target, to refresh participants’ memory (figure 1B). There were thus 104 trials in the pretests: 2×27 movement tests, 2×9 explicit tests, 4×8 movement practice trials.

In the subsequent *rotation practice* phase, participants performed 144 trials toward the practice direction with cursor movement being rotated relative to hand movement by 45°. The sign of the rotation switched between clockwise (CW) and counterclockwise (CCW) depending on the context condition, which here alternated every 8 trials. Before we first introduced the cursor rotation, we instructed participants that they would still control the cursor, but that the relation between the direction of hand and cursor movement would be changed and that this changed relation would be signaled by a red, instead of the already experienced green start circle.

Rotation practice was followed by a series of posttests to dissociate implicit, total and explicit learning. The *posttests* were structured like the pretests, except that the movement tests were repeated twice: the first repetition tested for implicit aftereffects by instructing participants to assume the rotation was switched off, reasoning that this would induce them to abandon potential aiming strategies and aim right at the target. The second repetition tested for total learning by instructing them that the rotation was switched on. For the explicit judgment tests, the rotation was instructed as switched on to test for explicit knowledge about the cursor rotation. Throughout the experiment, the presence or absence of the rotation was additionally cued by the color of the starting position (green = switched off, red = switched on), which participants were repeatedly reminded of.

### Specific experiments

The experiments differed in the type of contextual cue associated with the opposing cursor rotations. In all experiments, the rotation sign was associated with a visual workspace cue, meaning that start, cursor and target locations were presented with a constant x-axis shift of ¼ screen width (figure 1A). CW cursor rotation was always associated with display in the left half of the screen while CCW rotation was displayed in the right half. Hand movements were performed in a joint central workspace. We retained this for consistency with our previous experiments, where we found that it did not cue separate implicit memories and instead produced a pattern consistent with plan-based generalization (Hegele and Heuer 2010; Schween et al. 2018). Our main interest was thus on whether the added cues would enable separate implicit memories to be formed.

In *experiment 1*, the added cue was the visual identity of the cursor: participants either saw a hand icon or an arrow cursor. These cursor types were associated with the existing combination of visual workspace and cursor rotation in a way that was constant within, but counterbalanced across participants. As the cursor was visible once participants were in the vicinity of the start location, they could anticipate the upcoming rotation based on the cue in all movement trials. On explicit posttests, the cursor cue was attached to the far end of the ray that signaled participants response.

The added contextual cues for *experiment 2* were two different action effects: Participants were instructed that they would have to either “explode” the target or “paint” it. The effect that participants should intend was prompted by a screen message at the beginning of each block (German: “Zerstören!” or “Anmalen!”). Accordingly, an animated explosion or brushstroke appeared at the location where the cursor crossed target amplitude, accompanied by respective sounds. As in experiment 1, action effects were fixed to visual workspaces and rotation direction within, but counterbalanced across participants. During movement tests without feedback, participants received the onscreen message before each block and the audio was played to remind them of the intended action effects.

In *experiment 3*, the additional cue was the hand used to conduct the movement, where we always associated the left hand with the left visual workspace and the right hand with the right visual workspace. At the beginning of each block, participants were prompted about which hand to use by an onscreen message and we asked them to rest the idle hand in their lap.

### Data Analysis

We performed data analysis and visualization in Matlab (RRID:SCR_001622) and R (RRID:SCR_001905). X- and y-coordinates of the fingertip were tracked at 100Hz and low-pass filtered using MATLAB’s “filtfilt” command (4-th order Butterworth, 10 Hz cutoff frequency). We then calculated the movement-terminal hand direction as the angular deviation of the vector between start and hand location at target amplitude and the vector between start and target. We excluded the following percentages of trials for producing no discernible movement endpoints (usually because the trial was aborted): experiment 1: 5.3%, experiment 2: 4.4%, experiment 3: 3.6%, and an additional total of 24 trials for producing hand angles more than 120° from the ideal hand direction on a given trial. Explicit direction judgments were calculated as the deviation between the vector connecting the start position with the target and the participants’ verbally instructed direction judgement. To obtain our measures of aftereffects, total learning, and explicit judgments, we calculated the median of the three repetitions per target in each pre- and posttest, under each cue level, for each participant, and subtracted the individual median of pretests from their respective posttests to account for any biases (Ghilardi et al. 1995). As main outcome measure, we therefore report direction changes from pretest to the different posttests types, depending on test target direction and context cues.

### Statistical Analysis

As we were interested in whether or not the contextual cues enabled separate implicit memories, we focused on aftereffects and only report explicit judgments and total learning descriptively, for completeness. Furthermore, as generalization of explicit judgments appears to strongly depend on methodological details (Poh and Taylor 2018; Schween et al. 2018), we would not claim universal validity of our findings in this respect. In our main analysis, we aimed to infer whether implicit aftereffects assessed under each cue reflected only the cued transformation, or both, and if aftereffects differed depending on the cue level. We therefore fit two candidate functions to the group mean aftereffect data obtained under each cue, respectively. In line with our previous reasoning (Schween et al. 2018), we chose a single-peaked Gaussian to represent the hypothesis that aftereffects reflected only one learned transformation:

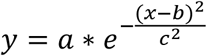

Here, *y* is the aftereffect at test direction *x*. Out of the free parameters, *a* is the gain, *b* the mean and *c* the standard deviation.

The hypothesis that aftereffects reflected two transformations was represented by the sum of two Gaussians:

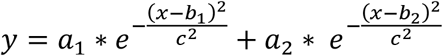

For this, we assumed separate gains *a*_1_; *a*_2_ and means *b*_1_; *b*_2_ but a joint standard deviation *c.* For fitting, we used Matlab’s “fmincon” to maximize the joint likelihood assuming independent Gaussian likelihood functions for the residuals. We restarted the fit 100 times from different values selected uniformly from the following constraints: −180° to 180° on *a*, 0° to 180° on b-parameters, 0° to 180° on c, and subsequently compared the fits with the highest likelihood for each model by Bayesian information criterion (BIC), calculated as:

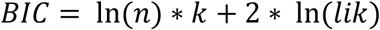

with *n* being the number of data points, *k* number of free parameters and *lik* the joint likelihood of the data under the best fit parameters.

In order to test if the generalization functions thus obtained differed significantly between the two context cues used in each experiment, respectively, we created 10000 bootstrap samples by selecting participants randomly, with replacement, and fitting on the across subject mean, starting from the best fit parameters of the original sample. For each sample, we calculated the difference between parameters obtained for each cue level. We considered parameters to differ significantly if the two-sided 95% confidence interval of these differences, calculated as the 2.5^th^ to 97.5^th^ percentile, did not include zero.

## Acknowledgements

We thank Manuela Henß, Simon Koch, Rebekka Rein, Simon Rosental, Kevin Roß and Vanessa Walter for supporting data collection.

## Author contributions

RS and MH designed experiments. LL conducted experiments. RS analyzed data and created figures. RS, LL, JAT and MH interpreted data. RS and LL wrote manuscript draft. RS, LL, JAT and MH revised manuscript and approved publication.

## Data availability

The datasets generated and analyzed during the current study are available from the corresponding author on reasonable request.

